# A Homology Guide for Pacific Salmon Genus *Oncorhynchus* resolves patterns of Duplicate Retention, Rediploidization and Local Adaptation Following the Salmonid Specific Whole Genome Duplication Event

**DOI:** 10.1101/2022.10.03.510689

**Authors:** Bradford Dimos, Michael Phelps

## Abstract

Salmonid fishes have emerged as a tractable model to study whole genome duplications (WGD) as this group has undergone four rounds of WGD, with a significant proportion of the genome yet to rediploidize. The fact that much of modern salmonid genomes retain duplicates from the most recent WGD while other regions have rediploidized creates complications for genetic studies by obscuring homology relationships and the necessity of filtering duplicate regions from many analyses. The difficulty this creates is particularly prominent in Pacific salmonids genus *Oncorhynchus* who are the focus of intense genetics-based conservation and management efforts owing to the important ecological and cultural role these fish play. To address this gap, we generated a homology guide for six species of *Oncorhynchus* with available genomes and used this guide to describe patterns of duplicate retention and rediploidization. Overall, we find that retained duplicates comprise over half of modern gene repertoires and that retained duplicate genes are enriched for genes involved in nuclear stability, while rediploidized genes which represent a smaller proportion of genes are heavily enriched in dosage sensitive processes such as mitochondria. Additionally, by reanalyzing published expression data from locally adapted strains of *O. mykiss* we demonstrate that retained duplicates are more likely to be associated with adaptive divergence than rediploidized genes, highlighting the potential of WGDs to promote adaptation. Finally, we demonstrate the utility of our homology guide by investigating the evolutionary relationship among genes highlighted as playing a role in salmonid life history traits or gene editing targets.

## Intro

Whole genome duplications (WGD)s have taken place many times in the history of eukaryotic evolution (Otto & Whitton, 2000). This includes vertebrates, whose common ancestor underwent two rounds of whole genome duplication referred to as the 2R hypothesis (Dehal & Boore, 2005). Some vertebrate taxa have undergone additional rounds of WGD, including a third round in teleost fish (Jaillon et al., 2004) and a fourth round in the common ancestor of salmonids termed the SS4R (Lien et al., 2016). Initially, WGDs events were studied in plants and thought to promote genomic instability and rearrangement (Pontes et al., 2004) leading to an evolutionary dead-end (Mayrose et al., 2011). However, work in animals does not support WGDs as a source of instability (Hufton et al., 2008), and evidence suggests that it may be a major driver of evolution and diversification (Soltis et al., 2014). In fact, WGDs can promote adaptation through the simultaneous duplication of all genes thereby creating large-scale opportunities for gene neofunctionalization and subfunctionalization (Force et al., 1999; Lynch, 2000; Ohno, Susumu, 1970). However, whether neofunctionalization or subfunctionalization predominate following a WGD have proved difficult to quantitatively address (Sandve, Rohlfs, & Hvidsten, 2018) as has the role WGDs play in promoting adaptation. Salmonid fishes have emerged as a tractable model to study WGDs as a substantial proportion of their genome has yet to rediploidize in the 80-106MYA since the SS4R (Gundappa et al., 2022; Lien et al., 2016).

Salmonid fishes including those in the genera *Salmo, Oncorhynchus, Hucho* and *Salvelinus* have provided insights into evolution following the SS4R as genome assemblies exists for many of these species (Christensen et al., 2018; Gao et al., 2021; Gundappa et al., 2022; Lien et al., 2016). It is believed that rediploidization in salmonids is on-going and proceeds in bursts. Under this model an initial wave of rediploidization occurred in the common ancestor of modern salmonids immediately following the WGD, followed by a period of relative stasis, then a smaller wave of lineage specific rediploidization (Gundappa et al., 2022). In salmonids many ohnologs (gene duplicates produced from a WGD event) have developed divergent expression patterns and thus are thought to be protected from rediploidization due to functional constraints (Gillard et al., 2021). However, whether this regulatory divergence occurs due to neofunctionalization or relaxed purifying selection remains an open question (Sandve et al., 2018). On the other hand, gene rediploidization appears to be due to structural rearrangements caused by transposable element insertions (Lien et al., 2016).

In addition to their importance in understanding genome evolution Salmonid fishes are also of tremendous cultural, economic and ecological significance. In North America, Pacific salmonid populations in the genus *Oncorhynchus* have provided food for indigenous communities since time immemorial (Atlas et al., 2021) and are a critical link for the flow of nutrients between the ocean and terrestrial ecosystems (Schindler et al., 2003). However, many populations of *Oncorhynchus* are experiencing precipitous declines (Crozier, Zabel, & Hamlet, 2007; Waldman & Quinn, 2022). These declines have prompted intense interest in understanding the genetic basis of ecologically important traits as well as the development of genome editing technology to aid in conservation efforts (Phelps, Seeb, & Seeb, 2020; Waples, Naish, & Primmer, 2020). However, the variation of gene copy number produced by differential rediploidization between species and inconsistent gene naming complicates these efforts (Limborg, Larson, Seeb, & Seeb, 2017; Rougemont et al., 2022) by making the identification of homologous genes, a prerequisite for either genetic analysis or genome editing efforts extremely challenging.

Given the substantial influence copy number variation can have on ecologically important traits and evolutionary patterns, we investigated gene duplicate retention and rediploidization across six Pacific salmon species of the genus *Oncorhynchus* with sequenced genomes for Chinook (*O. tshawytscha*), Coho (*O. kisutch*), Sockeye (*O. nerka*), Chum (*O. keta*), Pink (*O. gorbuscha*) and Rainbow trout (*O. mykiss*). For three species Chinook, Sockeye and Rainbow trout we investigated how gene duplicate retention/rediploidization has influenced genome structure. Additionally, we examined the potential of retained duplicates to facilitate adaption by reanalyzing gene expression from the redband rainbow trout model system (*O. mykiss gardeneri*) (Chen, Farrell, Matala, & Narum, 2018). Through our investigation we produced a gene family-based homology guide for *Oncoryhynchus* which includes both gene name (NCBI ID) and copy number that we hope we will serve as an easy to use resource for the salmonid community to address the issue of homolog identification. To demonstrate the utility of our homology guide we produce gene trees for gene families highlighted as influencing life-history variation or potential high value gene editing targets.

## Results and Discussion

### Gene Family Evolution Following the SS4R

In order to understand the process of duplicate retention and rediploidization in *Oncorhynchus* we grouped genes into homologous genes families and quantified the copy number for the six species with publicly available genomes at the time of analysis *O. tshawytscha, O. kisutch, O. mykiss, O. nerka, O. keta, O. gorbuscha* as well as Atlantic salmon (*Salmo salar*) and Northern Pike (*Esox Lucius*) as outgroups. While our method based on gene copy number is unable to differentiate between gene duplicates produced from the SS4R (ohnologs) versus gene duplicates which did not arise from the SS4R (paralogs) there is not yet a consensus best practice for determining ohnologs as methods such as gene tree-species tree reconciliations can have high false negative rates and exclude multi-copy gene families (Braasch et al., 2016; Inoue, Sato, Sinclair, Tsukamoto, & Nishida, 2015). Therefore, we chose to use a more inclusive strategy to identify putative ohnologs, which we term retained duplicates. Gene families were assigned into one of five categories: retained duplicates, rediploidizing, rediploidized, expanding or contracting based on comparison to *E. Lucius*. The number of genes families which are either retained duplicates, rediploidizing, rediploidized, expanding or contracting varied some by species but were relatively consistent with the exception of *O. keta* (Figure 1). The difference seen in *O. keta* were likely due reduced completeness and lower contiguity of this genome assembly as noted by the authors (Rondeau et al., 2021). Retained duplicates comprised between 51.7%-67.0% of gene families with only 27.5%-40.7% of families being rediploidized (Figure 1). Interestingly, we found that only about 2.13%-3.65% of homologous genes were in the process of rediploidizing. Expanding gene families made up between 2.50%-9.47% of all homologous gene, while contracting genes families comprised between 1.92%-2.90% of all homologous genes, indicating that while individual gene duplications and losses are occurring these processes are a smaller contributor to shaping the current gene repertoires of *Oncorhynchus spp*. than the SS4R. For a full breakdown of the percentage of gene families within each category see table 1.

**Figure 1:**
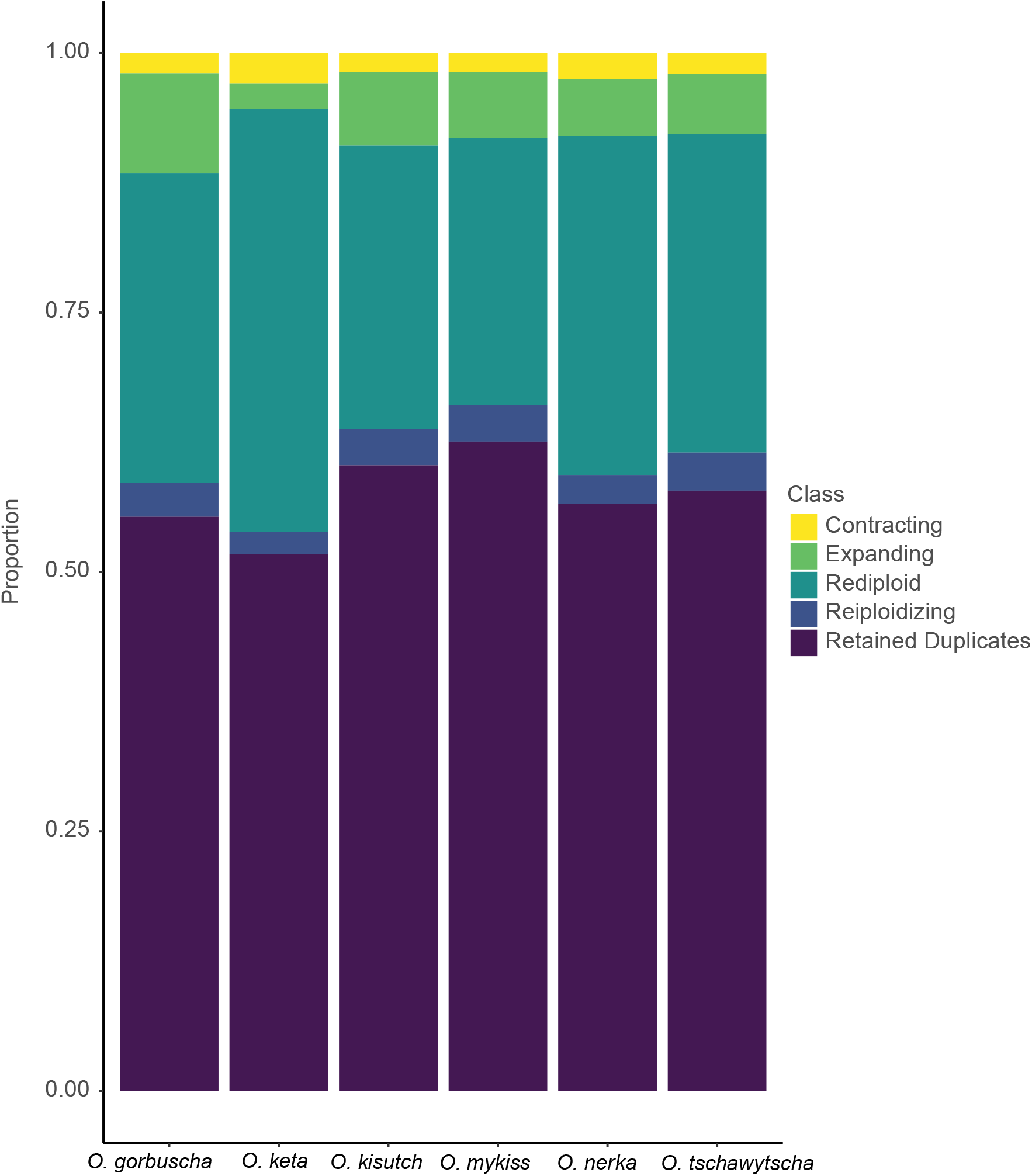
The relative proportion of the gene repertoire of each species belonging to each of the five gene duplication categories.

**Table 1.**

| Species | Contracting | Expanding | Rediploid | Reiploidizing | Retained Du | Total Genes |
| --- | --- | --- | --- | --- | --- | --- |
| *O_grobuscha* | 652 | 3220 | 10012 | 1092 | 18540 | 34016 |
| *O_keta* | 658 | 566 | 9208 | 482 | 11698 | 22612 |
| *O_kisutch* | 638 | 2421 | 9339 | 1203 | 20640 | 32975 |
| *O_mykiss* | 633 | 2238 | 8990 | 1222 | 21864 | 32667 |
| *O_nerka* | 764 | 1679 | 9969 | 853 | 17266 | 31700 |
| *O_tschwytsch* | 661 | 1927 | 10157 | 1215 | 19148 | 33452 |

### Retained Duplicates

We find widespread retention of duplicate genes across *Oncorhynchus*, comprising over half of the total gene repertoire of each species. The high number of retained duplicates is notable given that most gene duplicates are expected to be purged quickly (Innan & Kondrashov, 2010). However, if gene duplicates acquire new functions through sequence evolution or regulatory divergence (Force et al., 1999; Kondrashov, 2012) they can persist for millions of years. We observe high overlap in the number of the retained duplicates between species. Of the 12,579 gene families which are retained as duplicates in at least one species, 4,164 are retained universally across the six *Oncorhynchus* species, while species-specific duplicate retention was less common ranging from 150 in *O. keta* to 582 gene families in *O. mykiss* (Figure 2a).

**Figure 2:**
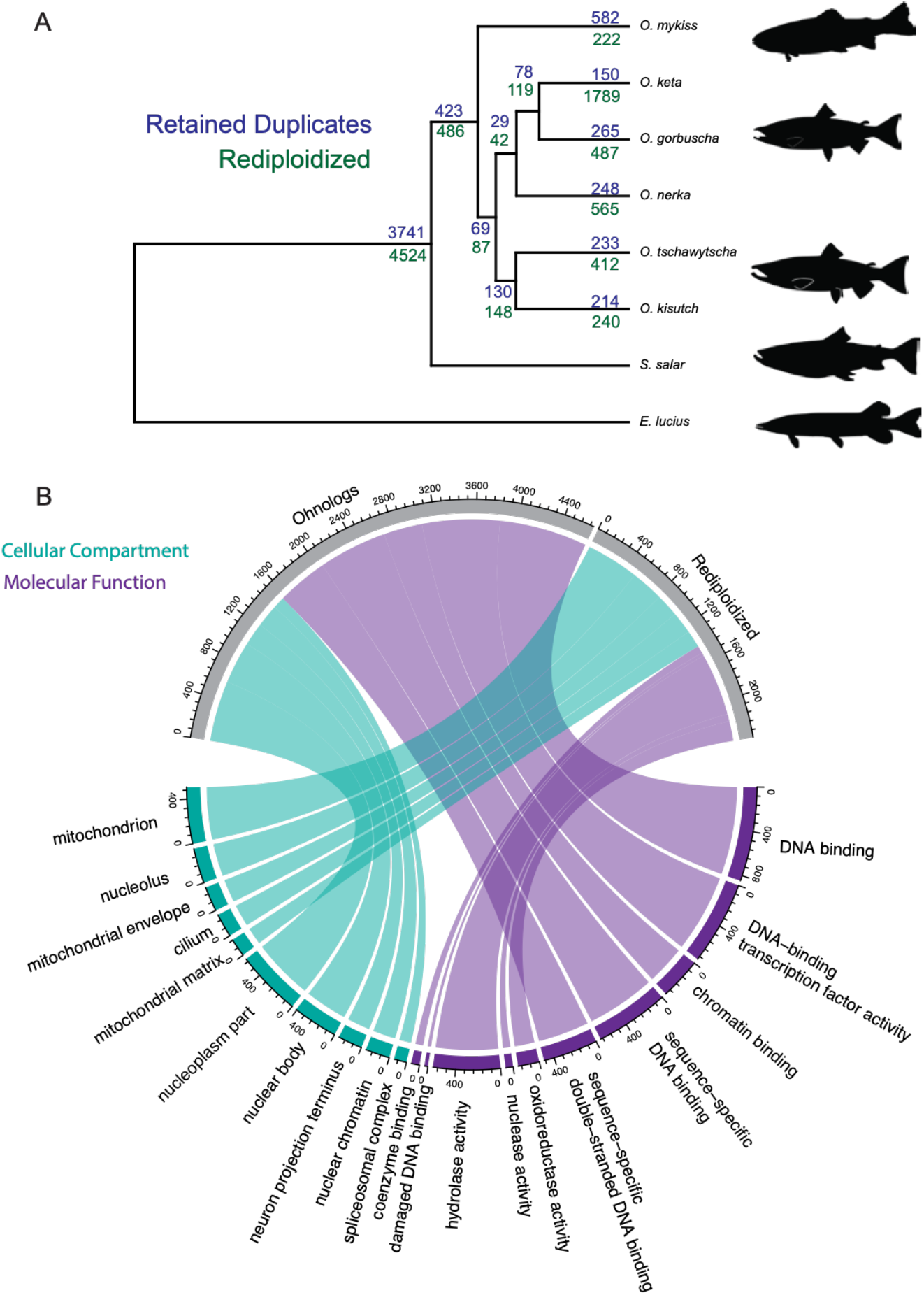
Ohnolog retention and rediploidizatioon. A) Species Tree with the number of genes families which are retained duplicates (blue) or have rediploidized (green). B) Chord diagram showing the visualization of select go terms, where GO terms are linked to either putative ohnologs or rediploidized gene families. Size of chord is proportional to the number of genes in a term which are either putative ohnologs or retained duplicates. Color denotes cellular compartment versus molecular function. For a full list of enrichments see Table S1.

As the number of universally retained duplicates is an order of magnitude larger than the number of retained duplicates in any one species, this indicates that a large proportion of salmonid genes (∼25%) are refractory to rediploidization, which could be due to constraints on gene function (Harper et al., 2021). To test this theory, we performed Gene Ontology (GO) enrichments on the universally retained duplicates. Our analysis revealed 15 cellular compartment enrichments which were heavily focused on the nucleus such as “nucleoplasm”, “catalytic step 2 splisosome” and “H4 acetyltransferase complex” (Figure 2b, Table S1). Likewise, these universally retained duplicates were also enriched for 29 molecular function terms heavily focused on DNA binding including, “chromatin binding”, “DNA binding” and “promoter specific chromatin binding”. However, there are also several other processes enriched among this list including, “acetyltransferase binding”, “peptide hormone binding” and “sphingolipid binding” (Figure 2b, Table S1). Given the essential role of these processes in basic cellular integrity and genomic stability our results raise the possibility that these genes may be maintained in a duplicate state due to their core functional significance. To this point the observed uniform copy number in all six salmonid species is unlikely to have been maintained across the 106 MYA (Gundappa et al., 2022) since the SS4R, unless the loss of one gene copy is highly selected against, supporting gene function as a potential driver of duplicate retention.

### Gene Rediploidization

Our findings identified a limited number of genes families that were actively in the process of rediploization. Taken together retained duplicates and fully rediploidized genes make up over 80% of all genes in the salmonid genome (Figure 1). This intriguing finding suggests that either the on-going wave of rediploidization (Gillard et al., 2021; Robertson et al., 2017) in *Oncorhynchus* may be nearly complete, or that the process of rediploidization can only be tolerated at a certain number of gene families simultaneously.

In salmonids rediploidization has been hypothesized to occur in bursts, where the first rediploidization event occurred in the common ancestor of salmonids, followed by a period of relative stasis, where another more recent round of rediploidization occurred in tandem with species diversification (Gundappa et al., 2022). Our results support this theory, since of the 14,776 total gene families which have rediploidized in at least one species, 5,010 have universally rediploidized in all six species (Figure 2a), while species-specific rediploiziation was less common, ranging from 222 in *O mykiss* to 1,789 in *O. keta* (Figure 2a). While the number of gene families which have rediploidized is larger than the number which have retained duplicates the copy number of the rediploizied gene families is lower and thus affects a smaller number of genes. The ancestral first wave of rediploidization, represented here as the universally rediploidized gene families (Figure 2a) affected a larger number of genes than the current wave of rediploidization, which is represented by the species-specific rediploidizations, similar to the findings of (Gundappa et al., 2022).

To determine what processes are affected by rediploidization we preformed GO enrichments on the universally rediploidized gene families. These enrichments analysis revealed 28 enriched cellular compartment terms, which relate to mitochondria and DNA including, “mitochondrion”, “mitochondrial matrix”, “DNA repair complex” and “replication fork” (Figure 2b, Table S1). The universally rediploidized genes were also enriched for 26 molecular function terms including “damaged DNA binding”, “oxidoreductase activity”, “tRNA binding” and “ligase activity” (Figure 2b, Table S1). Rediploidization of genes following a WGD is hypothesized to be driven by dosage balance constraints (Makino & McLysaght, 2010) and thus dosage sensitive genes would be expected to rediploidize to attain the gene copy number of the singleton (Woodhouse 2010). Taken together our data support the link between dosage sensitivity and rediploidization as the largest categories of enrichments are highly dosage sensitive processes such as mitochondrial metabolism (Toivonen et al., 2003). Mitochondria are particularly sensitive to gene dosage as mitochondrial activity requires the careful coordination of both the mitochondrial genome and the nuclear genome. Ensuring appropriate stoichiometric balance between the mitochondrial and nuclear genomes is critical for mitochondrial function (Wang et al., 2020). In fact, there are dedicated cellular pathways to ensure that both genomes work in harmony which are conserved across animal evolution (Dimos, Mahmud, Fuess, Mydlarz, & Pellegrino, 2019). WGDs thus have the potential to cause imbalance between nuclear and cytoplasmic genomes. Some polyploid plants solve the problem of stoichiometric balance through increasing the organellar genome copy number (Fernandes Gyorfy et al., 2021). However, our data indicate that the common ancestor of *Oncorhynchus* rediploidized mitochondrial genes in a manner consistent to achieve dosage balance between the nuclear and mitochondrial genome copy numbers. Interestingly, *S. salar* downregulates the expression of one copy of mitochondrial ohnologs (Gillard et al., 2021), thus, it appears that salmonids rediploidized a large number of nuclear encoded mitochondrial genes shortly after the SS4R and those that did not rediploidize developed compensatory expression patterns to avoid stoichiometric imbalance.

In addition to nuclear-encoded mitochondrial genes several other notable categories of genes, mostly related to nucleic acid repair are universally rediploidized across *Oncorhynchus* included those involved in DNA repair, replication fork, co-enzyme binding, tRNA binding and nuclease activity. While speculative, these processes might also be dosage sensitive and thus favor rediploidization, since they often involve the coordinated activity of many proteins. For example, tRNA copy number plays a role in determining codon usage (Duret, 2000) and processes such as DNA repair are sensitive to gene dosage (Chae et al., 2016). Overall, given the patterns observed both among the universally retained gene duplicates and the universally rediploidized gene families, functional constraints on genes appear to a major determinant of whether a gene family will rediploidize or be retained in its duplicate state.

### The Influence of Duplicate Retention and Rediploidization on Genome Structure

As rediploidization following a WGD is predicted to promote genomic instability (Semon & Wolfe, 2006) through structural rearrangements or transposable element insertion (Lien et al., 2016), we investigated if such a relationship can be observed in Pacific Salmon. To address this question, we identified syntenic regions between *O. mykiss, O. tshawytscha, O. nerka* and *S. salar* and tested if retained duplicates or rediploizied gene families were more or less likely to occur in syntenic regions. As rediploidization is expected to lead to genomic rearrangement (Sémon & Wolfe, 2007) and given that transposable element insertions are a major mechanism of rediploidization in Salmon (Lien et al., 2016) we would expect retained duplicates to be more likely to occur in more stable syntenic regions and rediploidized genes would be less likely to occur in these regions.

Overall, we observe high levels of genomic synteny as 38.9% of the *O. mykiss* genome, 49.2% of the *O. tshawytscha* genome and 48.2% of the *O. nerka* genome have obvious syntenic relationships with the *S. salar* genome (Figure 3a-c). Additionally, syntenic regions are found on every chromosome/linkage group with the exception of *O. tshawytscha* linkage group 11. While it is not apparent why *O. mykiss* has reduced levels of synteny compared to the other two species, two potential factors are that the *O. mykiss* genome used here (Arlee) was based on a clonal individual (Gao et al., 2021) or that the karyotype of *O. mykiss* is not fixed (Thorgaard, 1983). To exclude retained duplicates not arising from the SS4R we limited our search to universally retained duplicates which are likely to represent ohnologs. These putative ohnologs were more likely to occur in syntenic regions than a sized matched random sampling of genes in all three species (Figure 3d) (*O. tshawytscha* odds ratio 1.38492, p-value < 2.2e-16; *O. mykiss* odds ratio 1.406231, p-value < 2.2e-16; *O. nerka* odds ratio 1.516509, p-value < 2.2e-16; Fisher’s exact test). Additionally, rediploidized genes were less likely to occur in syntenic regions (Figure 3d) in *O. tshawytscha* (odds ratio 0.9212107, p-value = 0.002116; fisher’s exact test) and *O. nerka* (odds ratio 0.9208733 p-value = 0.00328; fisher’s exact test), but not *O. mykiss* (odds ratio 1.024598 p-value = 0.38267; fisher’s exact test) compared to a random sampling of genes.

**Figure 3:**
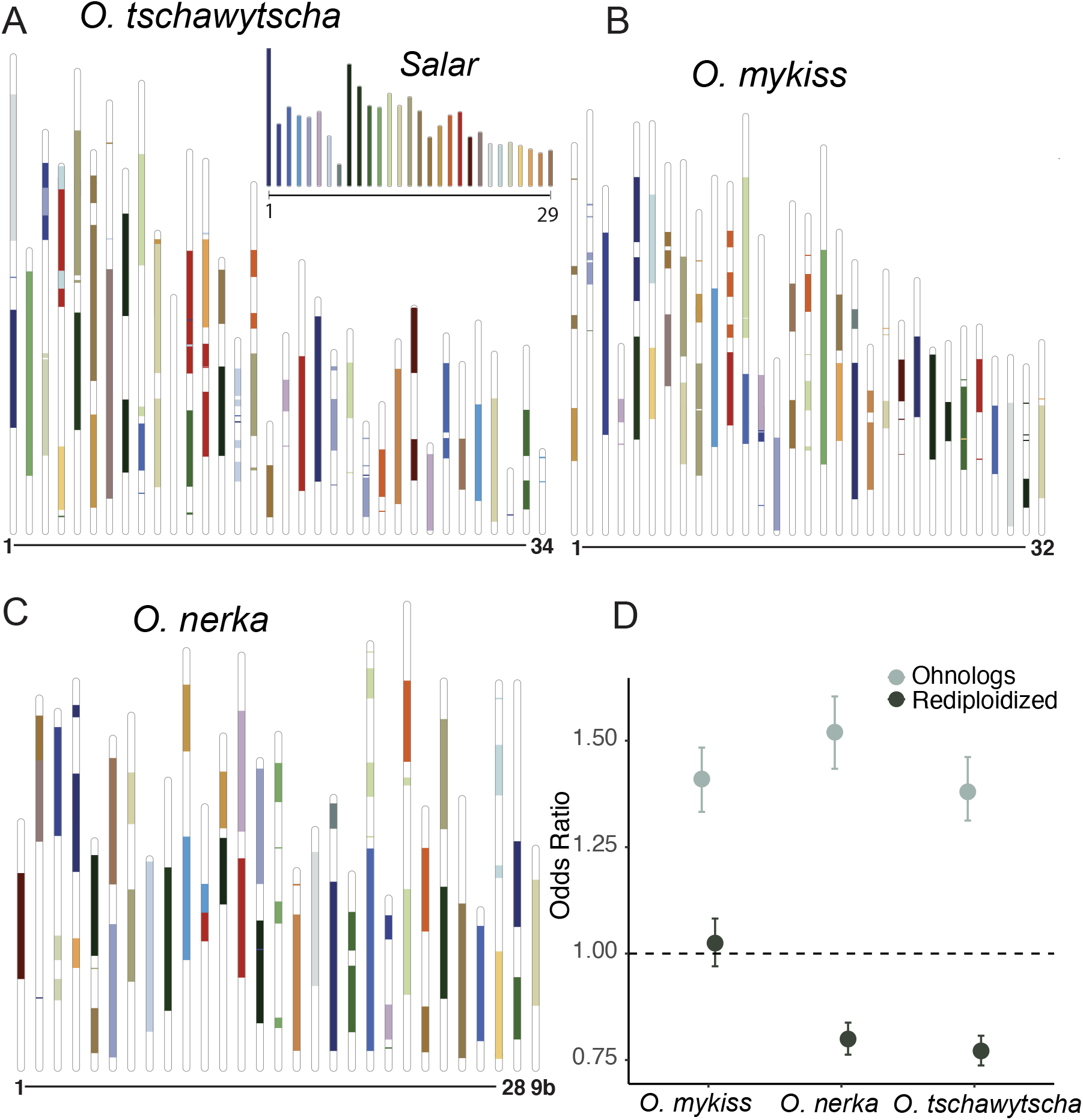
Synteny map for A) *O. tshwytscha*, B) *O. mykiss* and C) *O. nerka* compared to S. salar (inset). D) Odds ratio with 95% confidence intervals for the relationship between Ohnologs, Rediploidized genes and syntenic regions for each species.

We find general support for the relationship between genome synteny and ohnolog retention and rediploidization. Taken together with the observed gene function enrichment patterns, our data provide support for gene function as a likely determinant of whether a gene family will be retained as an ohnolog, or will rediplodize, which in turn may influence genomic structure. It should be noted however, that our analysis is unable to determine if gene retention/rediploidization causes genomic rearrangements or visa-versa. To this point if chromosomal rearrangements or TE insertions lead to loss of one the ohnolog copies then genomic rearrangement would be the driver of rediploidization. Supporting a role for gene function, genes which are involved in DNA stability seem to be particularly likely to be retained as ohnologs while genes involved in mitochondrial metabolism are more likely to have rediploidized. Interestingly, both retained duplicates and rediplodized genes which are shared between the six species of *Oncorhynchus* are far more numerous than those which are species specific. For the universally retained duplicates these genes are likely highly-refractory to copy number change as the copy number of these families have been maintained since the SS4R. For the universally rediploidized gene families the increased dosage following the SS4R was likely highly deleterious and organismal viability following the WGD was likely dependent on rapid rediploidization of these gene families.

### Retained Duplicates are Associated with Adaptive Divergence

Ohnologs have been widely suggested to promote adaptation, however only a few empirical examples exist which support this relationship in animals. To investigate this process, we analyzed RNAseq data taken from heart ventricles from three recently diverged populations of Redband Trout (*O. mykiss gairdneri*) which have adapted to live in cold montane, cool montane or warm desert streams (Narum & Campbell, 2015; Narum, Campbell, Kozfkay, & Meyer, 2010). To investigate the relationship between retained duplicates and adaptation, we used the evolutionary analysis of variance model to identify genes which have divergent expression patterns between populations. Of the 25,441 genes used in the model, 783 genes displayed divergent expression patterns. Of the retained duplicates 5.80% are divergently expressed between populations, which is a significantly larger proportion than the 2.95% of rediploidized genes which display divergent expression patterns (odds ratio 1.96645, p-value = 3.948e-14 Fischer’s exact test). This suggests that retained duplicates are more likely to be associated with local adaptation than rediploidized genes. Highlighting this, the retained duplicates with divergent expression patterns show clear differences in expression between the desert and cold montane populations, with cool montane populations showing intermediate expression profiles (Figure 4). The increased likelihood of retained duplications to demonstrate expression divergence also holds when considering universally retained ohnologs (odds ratio 1.814703, p-value = 2.108e-07 Fischer’s exact test). We found no evidence that our approach had difficultly differentiating paralogs due to low sequence divergence as the proportion of retained duplicates expressed above the minimum threshold was similar to the proportion of rediploidized genes. This finding provides empirical support that ohnologs can provide the raw genetic material to promote adaptative expression patterns.

**Figure 4:**
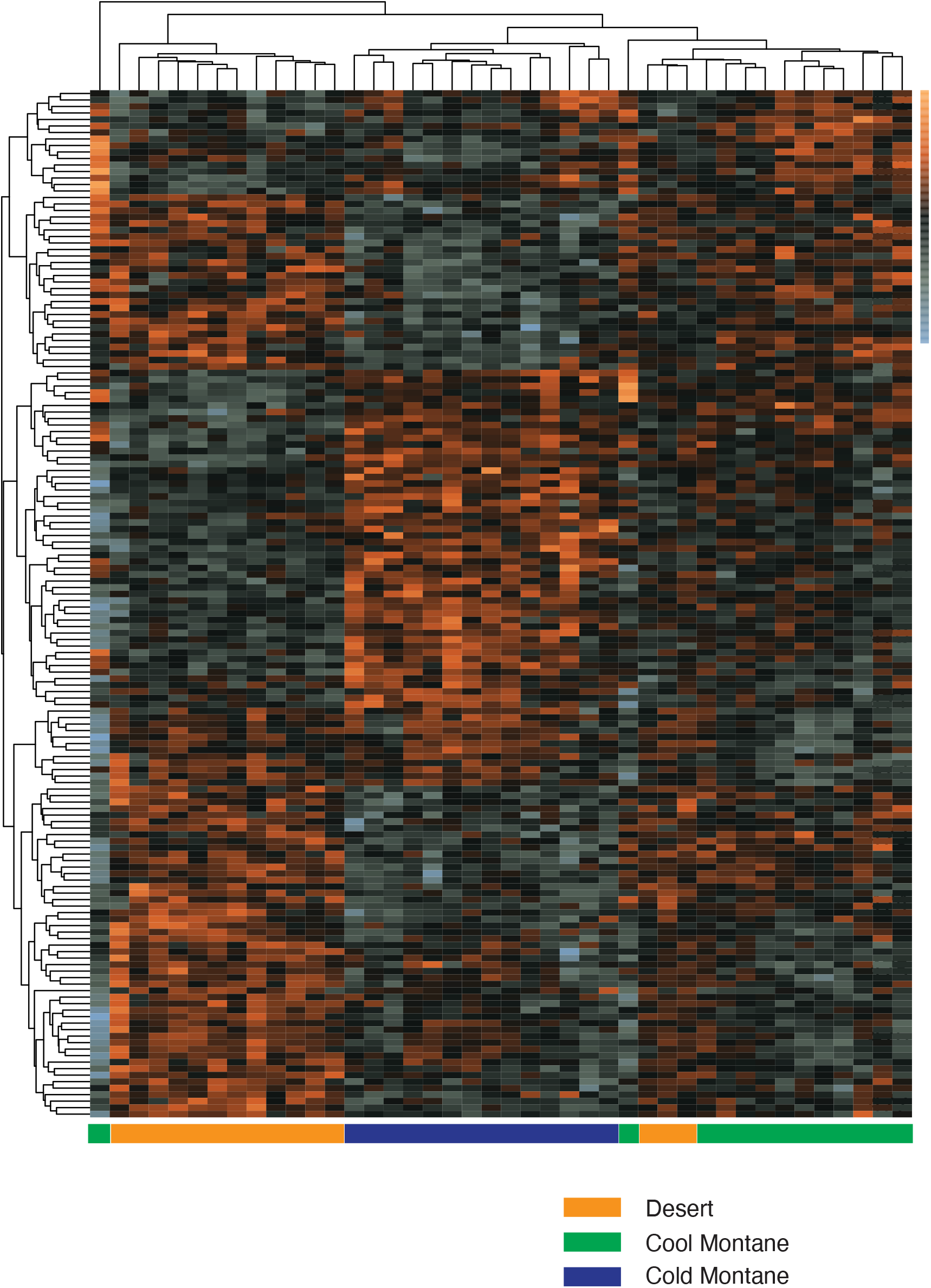
Retained duplicates demonstrating locally adaptive expression. Heatmap shows the z-score normalized expression of the 431 retained duplicates which show divergent expression profiles between cold montane, cool montane, and warm desert streams. Each column is a sample where population is denoted by color and each row is a gene. Orange gradient is increased level of gene expression whereas gray is decreased level of gene expression.

The recognition of WGDs as drivers of evolutionary adaptation is increasing (Glasauer & Neuhauss, 2014; Moriyama & Koshiba-Takeuchi, 2018) with the demonstration that some evolutionary novelties such as the development of electric organs in some fishes and the development of the bulbus arteriosus in zebrafish are due to functional divergence between ohnologs (Moriyama et al., 2016; Zakon, Lu, Zwickl, & Hillis, 2006). Recently, divergent expression of ohnolog copies was likewise demonstrated to promote adaption to salt water in mangrove plants (Xu et al., 2021). Our analysis of published data supports the role of WGDs in adaptive evolution by finding that retained duplicates are associated with locally adaptive expression patterns in trout hearts. It is important to note that this is based off of the gene expression profiles of a single tissue, and thus we are only observing a portion of locally adapted genes. However, as differences in cardiac performance have been shown to facilitate adaption to the differing thermal regimes experienced by these populations (Chen et al., 2018) the differing expression pattern in cardiac tissue is a good system to investigate genes involved in local adaptation. While it is not apparent if these expression patterns are due to neofunctionalization, subfunctionalization or altered gene dosage this underlies the utility of Salmonid fishes as a system for studying evolution following a WGD and provides support for predictions that WGDs can promote adaptation.

### Homology Guide Utility

While salmonids are an ideal model to study evolution following a WGD these fish are also intensely studied from a conservation and management perspective. Genetics is an essential component to modern salmonid management, however, variation in ploidy generated from the SS4R is difficult to incorporate (Limborg et al., 2017; Narum, Di Genova, Micheletti, & Maass, 2018), due to the paucity of data on gene copy number coupled with inconsistent naming schemes. Thus, we hope that our homology guide providing both the NCBI gene IDs and gene family size with common annotations across these six species of *Oncorhynchus* will prove a valuable resource for the Pacific salmonid research community. As an example, we demonstrate how this homology guide can be used to aid in the context of both genome-wide association studies and the emerging field of genome editing by illustrating how the SS4R has influenced the copy number and evolutionary relationships determined by Orthofinder2 of four previously highlighted gene families: *greb1l, vgll3, slc45a2*, and *dnd1*.

Run timing of coastal *O. mykiss* and *O. tshawytscha* in the western United States was shown to be nearly perfectly associated with variation in a genomic region containing the *greb1l* gene on chr. 28 (Hess, Zendt, Matala, & Narum, 2016; Prince et al., 2017; Thompson et al., 2020) and this gene has also been associated with age at maturity in *S. salar* (Cauwelier, Gilbey, Sampayo, Stradmeyer, & Middlemas, 2018). The gene *greb1l* is represented by gene family 5,379 and is double copy across *Oncorhynchus*, except for being single copy in *O. keta*, and triple copy in *S. salar* (Figure 5a). The *greb1l* gene tree represents standard ohnolog retention (except for the gene loss in *O. keta* and the extra duplication in *S. salar*), as each version of the ohnolog forms a monophyletic cluster. While the *greb1l* copy associated with run-timing is located on chr. 28 there is another *greb1l* copy located on chr. 10. This raises the question of the functional significance of the other *greb1l* copy and whether the ability of the *greb1l* copy on chr. 28 to determine run timing may be an example of neofunctionalization.

**Figure 5:**
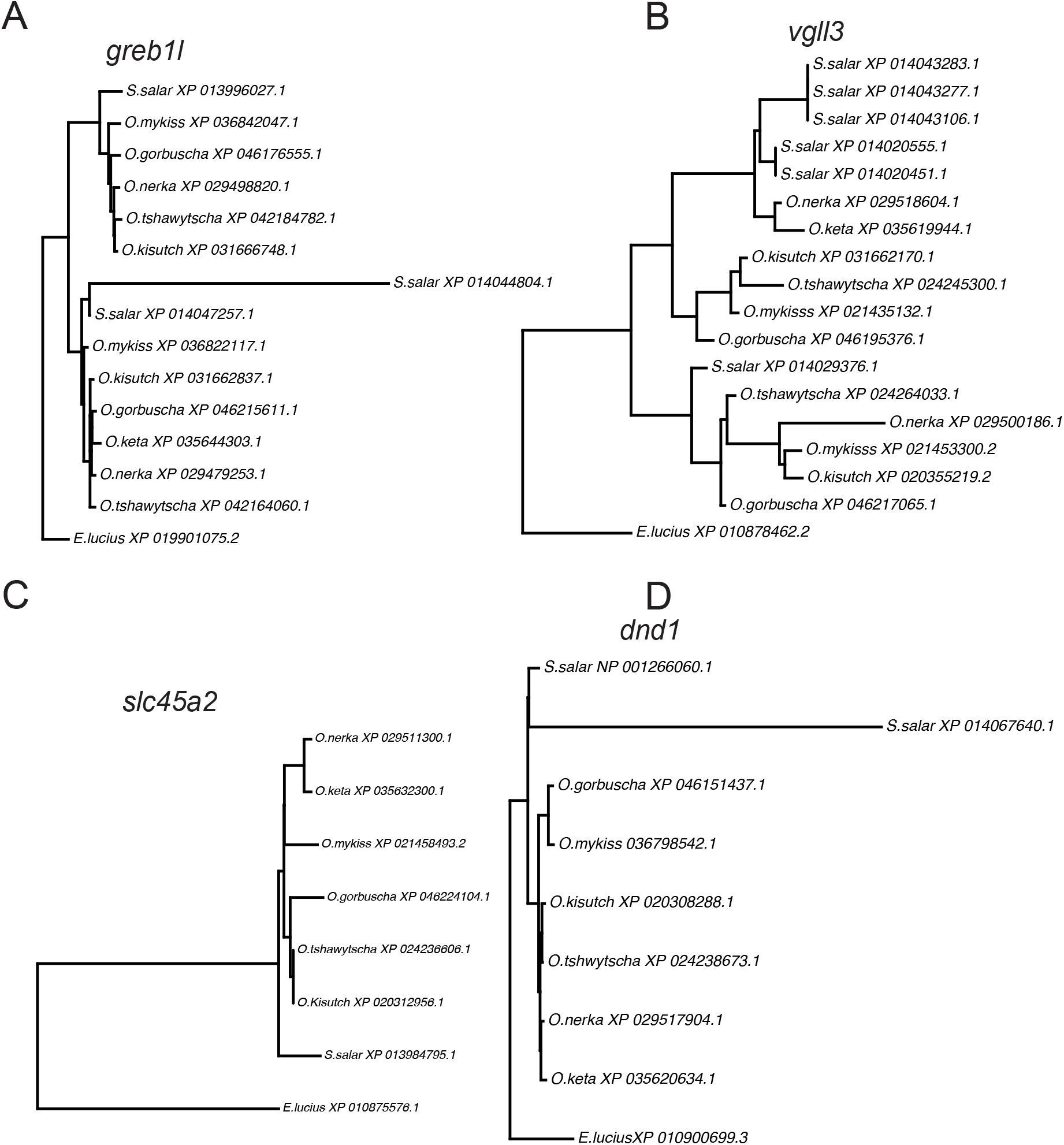
Gene trees of the genes highlighted as either being involved in life-history variation; (a) *grebl1*, and (b) *vgll3*, or as targets for gene editing; (c) *slc45a2* and (d) *dnd1*.

Age at maturity is an ecologically important life history trait and has recently been shown to be associated *vgll3* in *S. salar* (Ayllon et al., 2015). The gene *vgll3* is represented by gene family 1,099 and has the same copy number as *greb1l* with the exception of having six copies in *S. salar* (Figure 5b) and has an unclear evolutionary relationship likely due to the proliferation of the gene family in *S. salar*. Interestingly, the function of *vgll3* is not conserved between the two genera as *vgll3* has no influence on age at maturity in *Oncoryhunchus* (Waters et al., 2021) and thus the ability of *vgll3* to determine age at maturity may be an example of neofunctionlization of one of the duplicates in *S. salar*. This example underlies how gene copy number can influence function as well as the importance of incorporating ploidy in genome-based association studies for organisms which have experienced WGDs. As the field of ecological genomics continues to unravel more aspects of salmonid biology, we hope that our homology guide will prove to be a useful resource to translate findings across species.

A method gaining traction for functional genomics research or as a conservation tool is gene editing through the use of CRISPR technologies (Phelps et al., 2020). For CRIPSR technologies it is particularly important to understand gene copy number and homology. Without this information an edited animal may show no phenotypic differences due to the activity of a redundant gene copy, or alternatively a CRISPR targeting guide RNA may have multiple off-target effects if copy number is not accounted for. Salmonids have proven to be amenable to CRIPSR with the pigment transporter gene *slc45a2* commonly used as a marker gene to screen for an albino phenotype in edited fish (Edvardsen, Leininger, Kleppe, Skaftnesmo, & Wargelius, 2014; Straume et al., 2020). As an example of the utility of the homology guide, *slc45a2* represented by gene family 16,929 is single copy in all species (Figure 5c) and the gene tree of *slc45a2* is characteristic of a universally rediploidized gene. Thus, this gene will likely serve as an effective marker gene for editing in any species of *Oncorhynchus*. Another example is the dead-end gene (*dnd1)* which has been edited in *S. salar* and *O. mykiss* with the intent to produce fish with inherited sterility (Fujihara et al., 2022; Güralp et al., 2020; Wargelius et al., 2016). The gene *dnd1*, represented by gene family 13,707, is single copy across *Oncorhynchus* but is double copy in *S. salar* (Figure 5d). The high sequence divergence in one of the duplicates in *S. salar* is characteristic of rapid sequence evolution following duplication of *dnd1* which may suggest some functional divergence between the two retained duplicates. This may explain why targeting the one gene copy with lower sequence divergence is sufficient to block germ cell migration in these fish. As the field of genome editing in salmonids continues to grow we hope that this homology guide will become a useful tool for planning CRISPR targeting strategies.

## Conclusion

The relatively recent SS4R event makes salmonids an ideal system for studying the genomic consequences of WGD events. The SS4R has produced pronounced genetic variation, some of which has adaptive significance, which needs to be taken into account for practical applications of salmonid management and conservation. From an evolutionary prospective we demonstrate that gene function is major driver of whether ohnologs will be retained or will rediploidize, which either conserves or erodes chromosomal synteny, respectively. Additionally, we demonstrate that retained duplicates are more likely to develop divergent expression patterns than rediploidized genes. From a practical application perspective, we provide an easy to use homology guide with gene family name, gene copy number, and NCBI gene ID across *Oncorhynchus* and *S. salar*. As salmonid biology moves into the functional genomics age this study should provide a basis for how the SS4R has shaped the evolutionary history of these six species and should influence conservation strategies moving forward.

## Methods

### Homology and ohnolog identification

The gene repertoire of each species was identified through downloading the predicted proteome and genome annotation file from NCBI for *O. tshwytscha* (Otsh_v2.0) (Christensen et al., 2018; Narum et al., 2018), *O. mykiss* (USDA_OmykA_1.1) (Gao et al., 2021), *O. nerka* (O. ner_1.0) (Christensen et al., 2020), *O. keta* (Oket_V1) (Garvin, Saitoh, Churikov, Brykov, & Gharrett, 2010), *O. kisutch* (Okis_V2), *S. salar* (Ssal_v3.1), and *E. lucius* (fEsoLuc1.pri) (Ishiguro, Miya, & Nishida, 2003) in October 2021. *O. gorbuscha* (OgorEven_v1.0) (Christensen et al., 2021) was downloaded in Februrary 2022. Gene isoforms were removed based on transcriptional start site and only the longest isoform was retained. After filtering out isoforms we used Orthofinder2 (David M. Emms & Kelly, 2015, 2019) to assign genes families and construct gene trees. The species tree was constructed using the program species tree from all genes (STAG) (D.M. Emms & Kelly, 2018) as implemented within Orthofinder2. Genes families were classified based upon gene counts and comparison to *E. Lucius*. Gene families were classified as retained duplicates if the family was exactly twice as large in the focal species as in *E. Lucius*. Gene families which are undergoing rediploidization were larger in the focal species than *E. Lucius* but less than twice as large. Gene families which have rediploidized were exactly the same size in the focal species and in *E. Lucius*. Expanding gene families were more than twice as large in the focal species compared to *E. Lucius*. Contracting gene families were smaller in the focal species than they were in *E. Lucius*.

### GO enrichments

*S. salar* peptide sequences were annotated using the online portal of EggNOG mapper (Huerta-Cepas et al., 2017, 2019) using taxa auto-detection. *S. salar* annotations were used to assign gene names as well as GO terms to each gene family. GO enrichments were carried out for universally retained ohnologs across *Oncorhynchus* and universally rediploidized genes across *Oncorhynchus* separately. Enrichments were performed using the R script gene ontology with Mann-Whitney U test (GO_MWU) (Wright et al., 2017) using a fisher’s exact test with *S. salar* as the enrichment background. For the putative ohnolog enrichments, genes which were universally retained duplicates were given a score of 1 while all other genes were given a score of 0. For the rediploidized enrichments universally rediploidized genes were given a score of 1 while all other genes were given a score of 0. Identical parameters were used for each category (largest 0.25, smallest 50, clusterCutHeight = 0.25) and were kept consistent for both cellular compartment (CC) and molecular function (MF) enrichments.

### Synteny

Syntenic analysis was performed for three of the five species (*O. mykiss, O. tshawytscha* and *O. nerka*). Each species genome was initially separated by chromosome/linkage group and used as a query against the Atlantic salmon genome using LastZ (Harris, R. S., 2007) with parameters --chain --gapped --inner=1000. LastZ alignments were then chained together with axtchain with parameters -minScore=3000 -linearGap=medium, and then sorted using chainsort. Alignment chains were filtered with chainPreNet and then chained together with chainNet. Alignment nets were then made using netSyntenic. Nets were then converted into axt then maf format using netToAxt and axtToMaf respectively. Finally syntenic blocks were called using Maf2synteny with a minimum block size of 100kb (Kolmogorov et al., 2018). Syntenic blocks were then combined if consecutive blocks were in the same strand orientation and the gap between two consecutive blocks was less than 1 Mb. This process was then repeated for each chromosome. Genes were considered to be in syntenic regions if the start site of a gene fell within syntenic blocks on the chromosome/linkage group it was on based on the annotation file. The relationship between ohnologs/rediploidized genes and syntenic regions were quantified using Fischer’s exact test. All statistical analysis were carried in the R programming environment.

### Local Adaptation

The propensity of retained duplicates to be associated with local adaptation was tested by reexamining RNAseq data from the Redband Trout system (Chen et al., 2018). Raw reads were downloaded from the NCBI short read archive SRP109007 and F1 hybrids were excluded from further analysis. Adaptors and low-quality reads were removed with TrimGalore (Felix Krueger, 2021). These filtered reads were then quantified using Salmon (Patro, Duggal, Love, Irizarry, & Kingsford, 2017) with an index size of 31, decoys were constructed using genome sequences and individual quantification files were combined using TXimport (Soneson, Love, & Robinson, 2015). The read count table was then vst normalized in Deseq2 (Love, Huber, & Anders, 2014) by population after removing transcripts with less than an average of 10 counts. Our generated count matrix and the phylogenetic tree produced from these populations based on whole genome resequencing data (Chen & Narum, 2021) were used in the Expression and Analysis of Variance model (Rohlfs & Nielsen, 2015). This framework models expression level of a gene as a trait that evolves across a phylogeny using an Ornstein–Uhlenbeck process to determine a trait optima. The variance from this optima both between and within populations is then compared and significant variation is determined by a likelihood ratio test using chi-square distribution with one degree of freedom. The proportion of retained duplicate versus repdiploidized genes which are locally adapted was compared with a Fisher’s exact test.

## Data Accessibility

All data and code need to replicate the results of this study are available on github: https://github.com/braddimo/Oncorhynchus_Ohnologs

## Acknowledgments

This material is based upon work supported by the NSF Postdoctoral Research Fellowships in Biology Program under Grant No. 2109355.

## Author Contributions

Writing and Conceptualization: BD and MP. Data analysis: BD.

